# Leveraging protein language and structural models for early prediction of antibodies with fast clearance

**DOI:** 10.1101/2024.06.08.597997

**Authors:** Parisa Mazrooei, Daniel O’Neil, Saeed Izadi, Bingyuan Chen, Saroja Ramanujan

## Abstract

Monoclonal antibodies (mAbs) with long systemic persistence are widely used as therapeutics. However, antibodies with atypically fast clearance require more dosing, limiting their clinical usefulness. Deep learning can facilitate using sequence-based modeling to predict potential pharmacokinetic (PK) liabilities before antibody generation. Assembling a dataset of 103 mAbs with measured nonspecific clearance in cynomolgus monkeys (cyno), and using transfer learning from large protein language models, we developed multiple machine learning models to predict mAb clearance as fast/slow clearing. Focusing on minimizing misclassification of potentially promising molecules as fast clearing, our results show that using physicochemical properties yielded up to 73.1+/-1.1% classification accuracy on hold-out test data (precision 65.2+/-2.3%). Using only sequence-based features from deep learning protein language models yielded a comparable performance of 71+/-1.4% (precision 65.5+/-2.5%). Combining structural and deep learning derived features yielded a similar accuracy of 73.9+/-1.1%, and slightly improved precision (68.3+/-2.4%). Features important for classifying fast/slow clearance point to charge, moment, and surface area properties at pH 7.4 as well as deep learning derived features. These results suggest that the protein language models provide comparable information and predictive performance of clearance as physicochemical features. This work provides a foundation for in silico prediction of protein pharmacokinetics to inform antibody candidate generation and early deprioritization of designs with high risk of fast clearance. More generally, it illustrates the value of transfer learning-based application of protein language models to address characteristics of importance for protein therapeutics.

## Introduction

The development, approval, and clinical use of monoclonal antibody (mAb) therapeutics has grown substantially over the past 2 decades. As of January 2023, 145 mAbs were approved or in review in the EU or US [1]. Unlike their small molecule counterparts, mAbs are typically administered intravenously or subcutaneously to avoid degradation. However, the long systemic persistence (long half-life, low clearance) of mAbs, due largely to recycling via the neonatal Fc receptor (FcRn), enables less frequent administration compared to small molecule drugs. Antibodies that exhibit unfavorable physicochemical properties can exhibit in vivo aggregation, atypical distribution, nonspecific/off-target binding, or other behaviors that accelerate clearance and limit developability and clinical utility.

Even for a given target, mAbs with similar isotype and backbone can show significant differences in clearance in human, and in cynomolgus monkey (cyno), the preferred species for preclinical PK studies. In order to identify lead candidate antibodies in preclinical R&D, in vitro assays as well as in silico approaches have been previously developed that show association with non-specific clearance measurements in cynos [2,3]. Hotzel et al. introduced an assay based on ELISA detection of non-specific binding to baculovirus particles that could identify mAbs with higher risk of fast clearance [2]. While assays such as this are currently incorporated into workflows for therapeutic antibody screening and lead generation, they can only be performed after generation of the molecules, are time consuming and still suboptimal predictors. Based on an in silico analysis of a dataset of 61 mAbs, Sharma et al. previously showed that mAbs with faster clearance in cynos tend to have either higher hydrophobicity in complementarity determining regions (CDRs) of their variable domain (Fv), or extreme Fv charge [3]. Although these findings were based on linear correlation analyses, they suggest promise for combining structural properties in more sophisticated in silico models to predict mAb clearance.

Traditionally, protein structure and physicochemical properties have been calculated from sequence using molecular models based on structure, energetics calculations and homology with other proteins [16, 11]. Associations with descriptive features or rules from structural modeling have been used individually or in combination to predict protein properties of interest [3, 13]. Recent technological advancements have massively increased the number of protein sequences available in publicly accessible databases. This data along with significant advances in machine learning (ML) and specifically natural language processing have driven impressive progress in sequence-based prediction of protein structure and binding [14,15]. Well known examples of such models are AlphaFold and RoseTTAFold which have achieved remarkable accuracy in prediction of protein structures and interactions [14,15].

Neural networks are also being applied to the design of proteins and optimization of sequences toward desired functions [5,8,10]. However, centralized datasets containing large numbers of monoclonal antibodies with measured developability properties such as clearance are still lacking, limiting the ability to apply these methods for prediction of specific mAb properties. We can instead exploit advances in natural language models, unsupervised learning, and transfer learning to extract protein representations from task-agnostic protein language models trained on millions of protein sequences. These models can then be fine-tuned on datasets of antibodies [10, 18] and learned information can be then transferred to mAbs. During this process, the structural information contained within the sequence is converted into a multi-scale organization vector that semantically incorporates the same information. These vectors contain a range of information from residue-level features to features related to the homology of proteins [5, 8]. Such vectors are called *embeddings* and can be used by secondary machine learning methods to improve prediction of biological properties [6, 18]. Since the embeddings are so information rich, fewer training examples are needed for property prediction, allowing for use of smaller data sets. An example of such transfer learning is shown by Harmalkar et al. to predict antibody thermostability [18].

Using machine learning approaches, we have developed multiple in silico models for predictive classification of mAbs with fast versus slow clearance in cyno, based only on the mAb sequence. In this work, we explore machine learning models using: (1) physicochemical features extracted from molecular structural modeling and (2) embeddings based on protein language models. We explore the relative performance of the two approaches and assess potential benefit in combining both sets of features with the goal of maximizing prediction accuracy and precision in order to avoid incorrectly classifying potentially promising molecules as fast clearing. This work provides a proof of concept for sequence-based prediction of in vivo PK, laying the groundwork for use of transfer and deep learning in early sequence-based PK liability screening and for future methodological improvements.

## Results

### Characterization of monoclonal antibodies

Pharmacokinetic data were compiled for a set of 103 conventional monoclonal antibodies previously tested in preclinical development studies at Genentech. 102 of these are IgG1 isotypes and 1 is IgG4. The dataset covers a wide range of non-specific clearance values as measured in cynomolgus monkeys (1.6 - 249 mL/day/kg) (Figure 1). We considered 8 mL/day/kg as the cut-off to differentiate slow versus fast clearing mAbs based on internal observations as well as published analysis by Yadav et al. [19]. Of 103 mAbs, 62 (60.2%) had slow clearance (<=8 mL/day/kg) while 41 (39.8%) of mAbs had fast clearance (>8 mL/day/kg). Previously, Hotzel et. al investigated the association between synthetic library antibodies and fast clearance and found insufficient evidence for such an association [Hotzel et. al]. Our data on 103 mAbs also did not provide sufficient evidence of association between the variable domain groups and fast/slow clearance of antibodies (Figure 1 and Supp. Figure 1).

**Figure 1.**
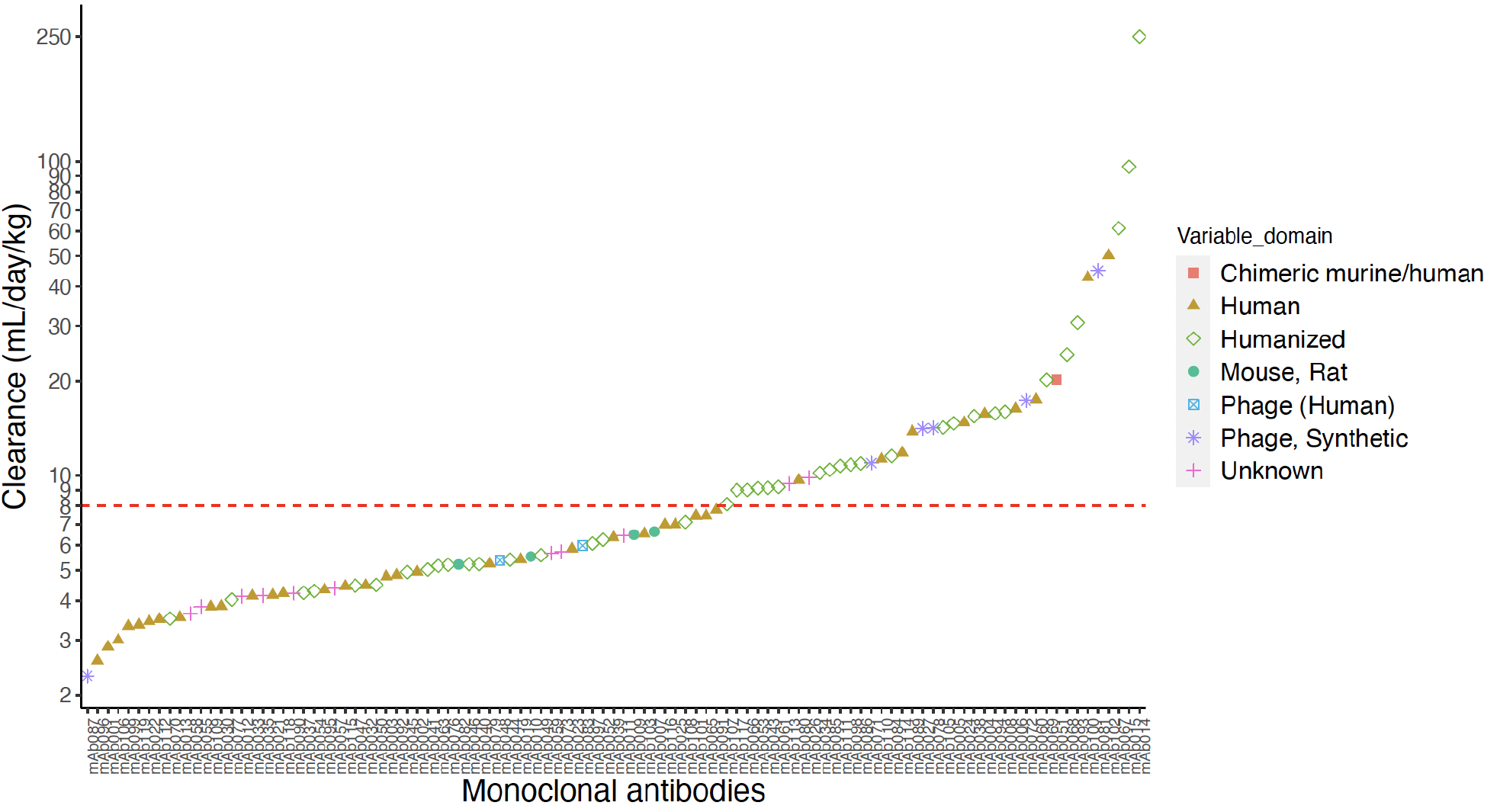
Mean clearance values of antibodies (n=103) show a wide range in cynomolgus monkeys. Variable domain information for antibodies are shown as follows: Chimeric murine/human (square), Human (triangle), Human synthetic phage (diamond), Humanized (circle), Mouse/Rat (square crossed), Phage/Human (star), Phage/Synthetic (plus). Variable domain information for 11 mAbs were not available (circle crossed).

### Structure-based features to represent physicochemical properties of mAbs

Certain physicochemical properties of mAbs have been previously shown to predict desirable features for developability, such as low/acceptable viscosity at high concentration [13]. Here, we use protein structure features of the Fab domains and electrostatic multipole moments and charge features of full-length antibodies to inform our machine learning models (Figure 2). To this end, protein properties for each Fab region were calculated using the Ensemble pH sampling module available in Molecular Operating Environment (MOE) 2019 [11] (see Materials and Methods for greater detail).

**Figure 2.**
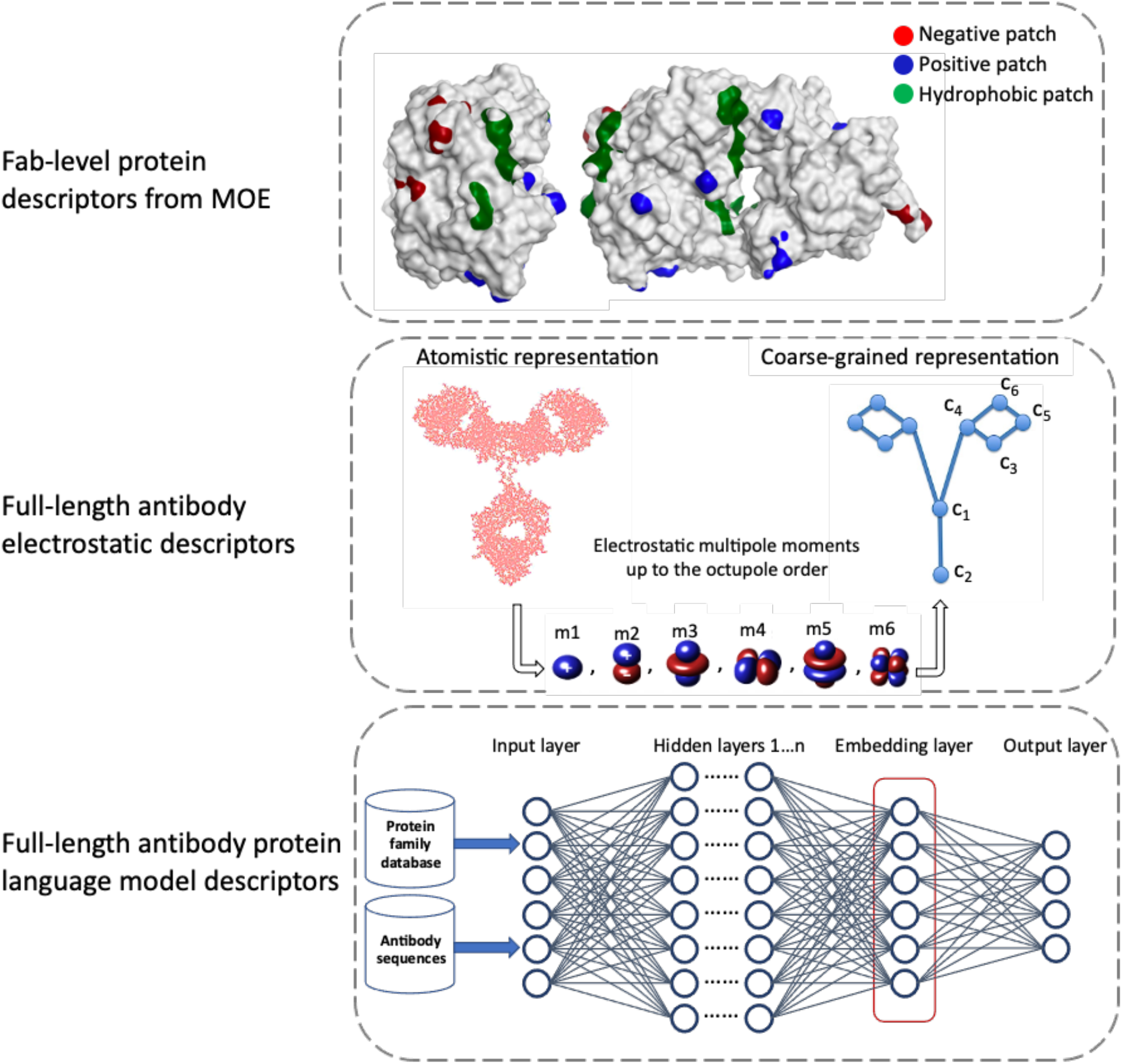
Schematic representation of descriptors; note that the schematic of the protein language model shown here is illustrative and the actual architecture can vary.

In addition to MOE protein properties calculations, we also calculated the low-order molecular multipole moments in the expansion of the full-length antibody charge distributions. The calculated moments are the monopole (m1), dipole (m2), linear and square components of quadrupole (m3 and m4), and linear and square components of octupole moments (m5 and m6). We also calculated six independent coarse-grained (CG) charges (c1, c2, c3, c4, c5, c6) that optimally reproduce these low-order molecular multipole moments [13](Figure 2). The details of the structure-based feature calculations are also provided in the Materials and Methods section.

The combination of MOE protein properties and the multipole moment and CG charges result in 33 unique physicochemical features which can be calculated at any specified pH. Among 33 features at pH 7.4, we observed significant association between only 4 computed protein properties and clearance of the mAbs (Wilcoxon test, p-value <0.05) (Supp. Figure 2), in line with what has been reported by Sharma et al. [3]. Although these features were associated with fast/slow clearance, none can be used as a predictive parameter alone. To assess whether different pH levels provide any additional information for the machine learning models, we considered two sets of descriptors: the 33 physicochemical features calculated at pH 5.5, and those same features calculated at Ph 7.4. The two pH levels correspond to systemic physiological pH (7.4) and the acidic environment found in the intracellular granules involved in FcRn-mediated antibody recycling. A complete list of features is provided in Supplementary table 1.

### Feature engineering improves model performance

We first developed and assessed the performance of classical machine learning models based on the physicochemical properties for identification of fast clearing mAbs. We used AutoML to find the optimal model architecture. Models were created with the feature sets at the two pHs. Table 1 lists the different feature sets included. To compensate for the small size of our dataset and avoid overfitting, we employed Monte Carlo cross validation for performance evaluation. We performed 100 iterations with different train/test splits of data. In each iteration, 103 molecules were randomly assigned into training and test sets based on a 80%-20% split of the dataset respectively. The mean and standard deviation of the model performance on test data based on different metrics were then computed and reported over the hundred iterations. Figure 3 shows the performance of classical machine learning models for pH 5.5 and pH 7.4. Over all the computed metrics, the pH 7.4 model performed better. Indeed, the low pH model performs comparably to a random, non-informative model.

**Table 1.**
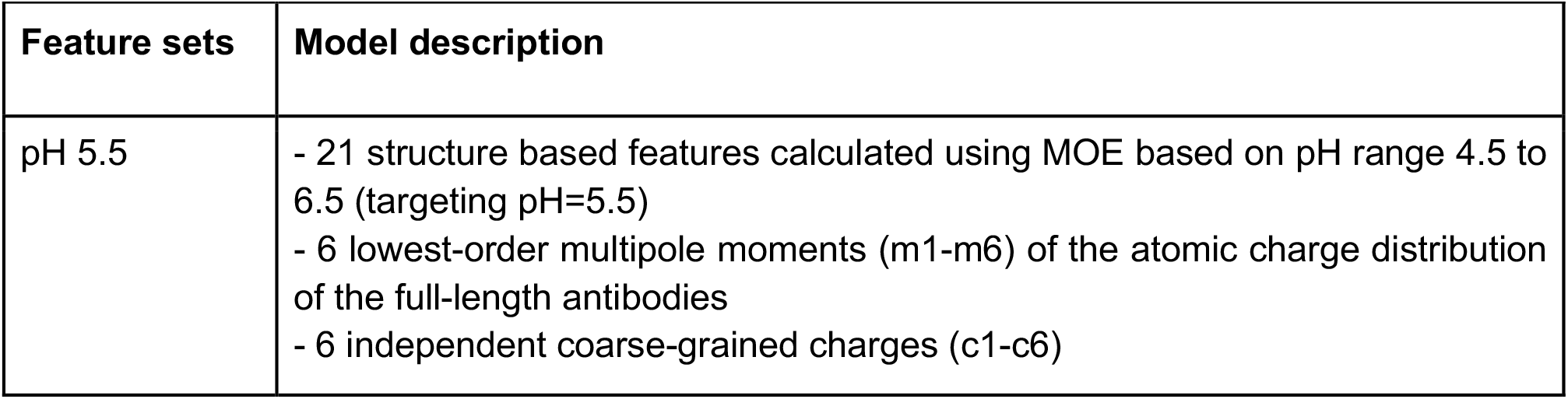
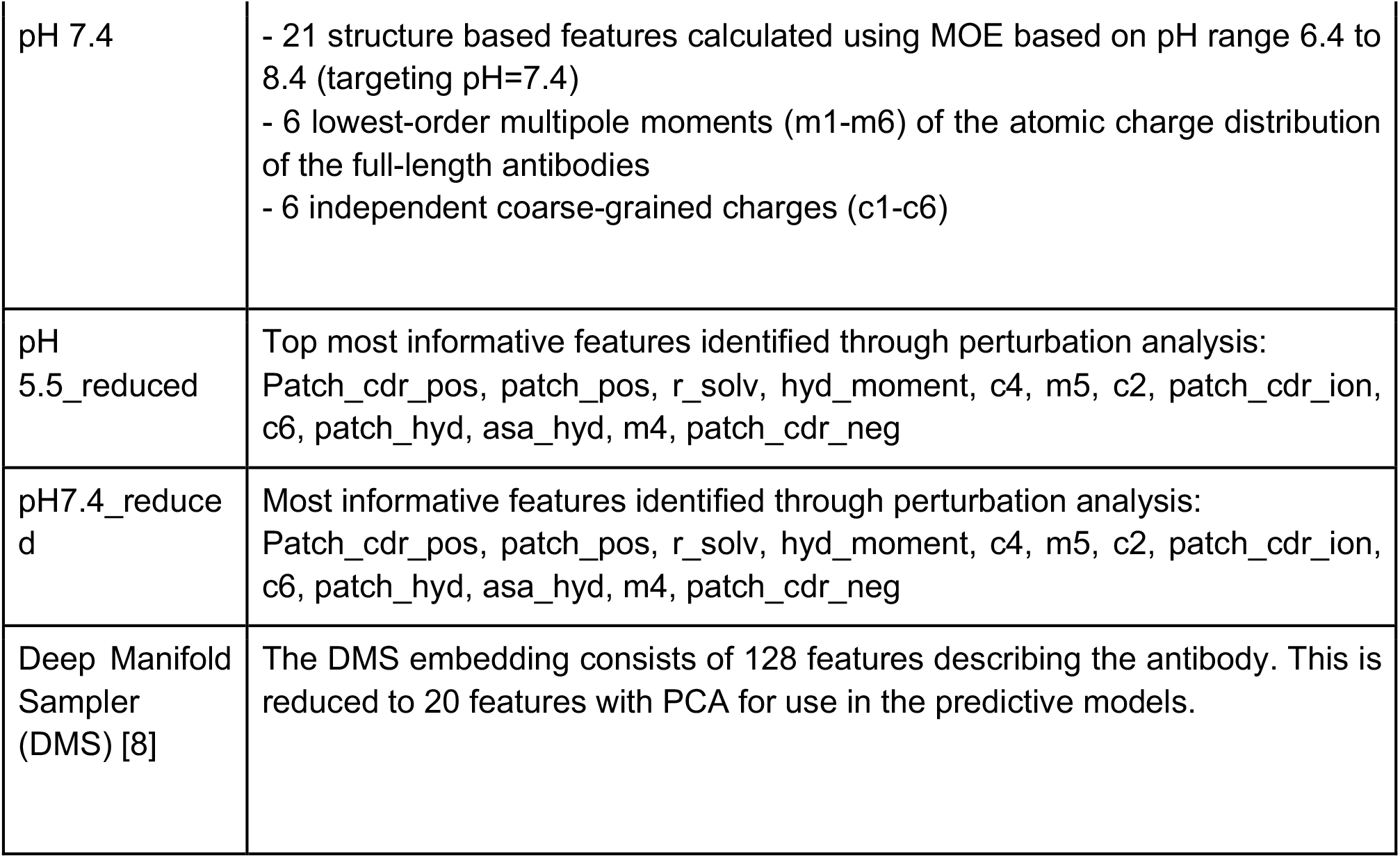
[18] Different methods used and their parameters and feature description. For further details of each model please refer to the Materials and Methods section. Note that “reduced” in feature sets’ names refers to the set of features obtained based on feature importance permutation analysis.

| Feature sets | Model description |
| --- | --- |
| pH 5.5 | - 21 structure based features calculated using MOE based on pH range 4.5 to 6.5 (targeting pH=5.5)<br>- 6 lowest-order multipole moments (m1-m6) of the atomic charge distribution of the full-length antibodies<br>- 6 independent coarse-grained charges (c1-c6) |
| pH 7.4 | - 21 structure based features calculated using MOE based on pH range 6.4 to 8.4 (targeting pH=7.4)<br>- 6 lowest-order multipole moments (m1-m6) of the atomic charge distribution of the full-length antibodies<br>- 6 independent coarse-grained charges (c1-c6) |
| pH 5.5_reduced | Top most informative features identified through perturbation analysis: Patch_cdr_pos, patch_pos, r_solv, hyd_moment, c4, m5, c2, patch_cdr_ion, c6, patch_hyd, asa_hyd, m4, patch_cdr_neg |
| pH7.4_reduced | Most informative features identified through perturbation analysis: Patch_cdr_pos, patch_pos, r_solv, hyd_moment, c4, m5, c2, patch_cdr_ion, c6, patch_hyd, asa_hyd, m4, patch_cdr_neg |
| Deep Manifold Sampler (DMS) [8] | The DMS embedding consists of 128 features describing the antibody. This is reduced to 20 features with PCA for use in the predictive models. |

**Figure 3.**
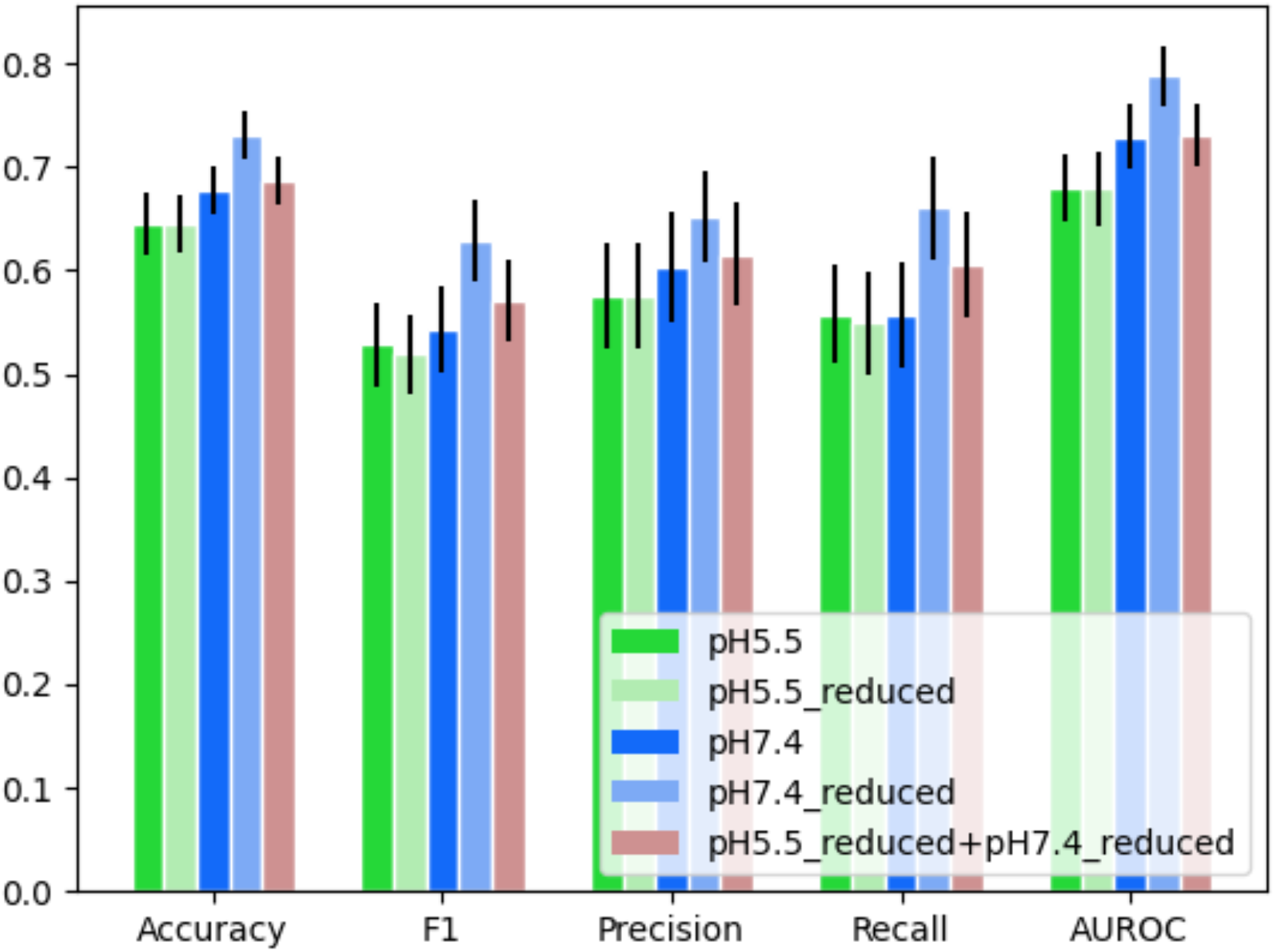
Performance evaluation and comparison of different feature sets. The pH 5.5 feature sets underperform the pH 7.4 feature sets. Using a set with a limited number of features improves all metrics for the pH 7.4 model. Black lines indicate the standard deviation. Note that “reduced” in model names refers to models trained on reduced number of features based on feature importance permutation analysis.

Since our analysis revealed a large number of highly correlated descriptors, (Supp. Figure 3), we sought to assess the effect of reducing the number of descriptors on the performance of the machine learning models. Feature importance permutation analysis was used to identify the top most informative features for the pH 5.5 and pH 7.4 feature sets (labeled as pH5.5_reduced and pH7.4_reduced respectively)(Table 1). As shown in Figure 3, reducing feature space improved the performance of the pH 7.4 feature set but had no effect on the low pH counterpart. This analysis supports the idea that when working with a small dataset and large number of descriptors, feature reduction by selecting the most relevant features can improve the model performance, but only when the features are inherently informative [21, 22]. Of the machine learning models based on physicochemical features, the model based on the pH7.4_reduced feature set performed best with an accuracy of 73.1±1.1% and a precision of 65.2±2.3%. We next sought to assess the performance of an objective, automated approach as opposed to a knowledge-based feature engineering.

### Model development incorporating embeddings learnt from a pre-trained protein language model

We next explored machine learning algorithms leveraging protein language model embeddings. The overall model structure of our ML pipeline that exploits embeddings from pre-trained protein language models is shown in Figure 4. The first stage of the ML pipeline is a deep neural network which produces embeddings of the mAbs. Deep Manifold Sampler (DMS) is a transformer-like model pre-trained on 20 million protein sequences from the protein family database, Pfam[8], and further fine-tuned on a dataset of 100K paired sequences of antibodies from Observed Antibody Space (OAS) [9, 10]. This model yields a 256 feature embedding for the entire antibody molecule. Principal component analysis was used to reduce the feature set to 20 features in total. The second stage in the pipeline is a Gaussian Naive Bayes classifier which leverages the embeddings produced in the first stage to predict the clearance. For more details of the model architecture, please refer to the Materials and Methods section. This PK prediction model based on the protein language model embeddings achieved a performance of 71±1.4% accuracy and 65.5±2.5% precision, comparable to that of the structure-based models using the pH7.4_reduced set alone (Figure 5).

**Figure 4.**
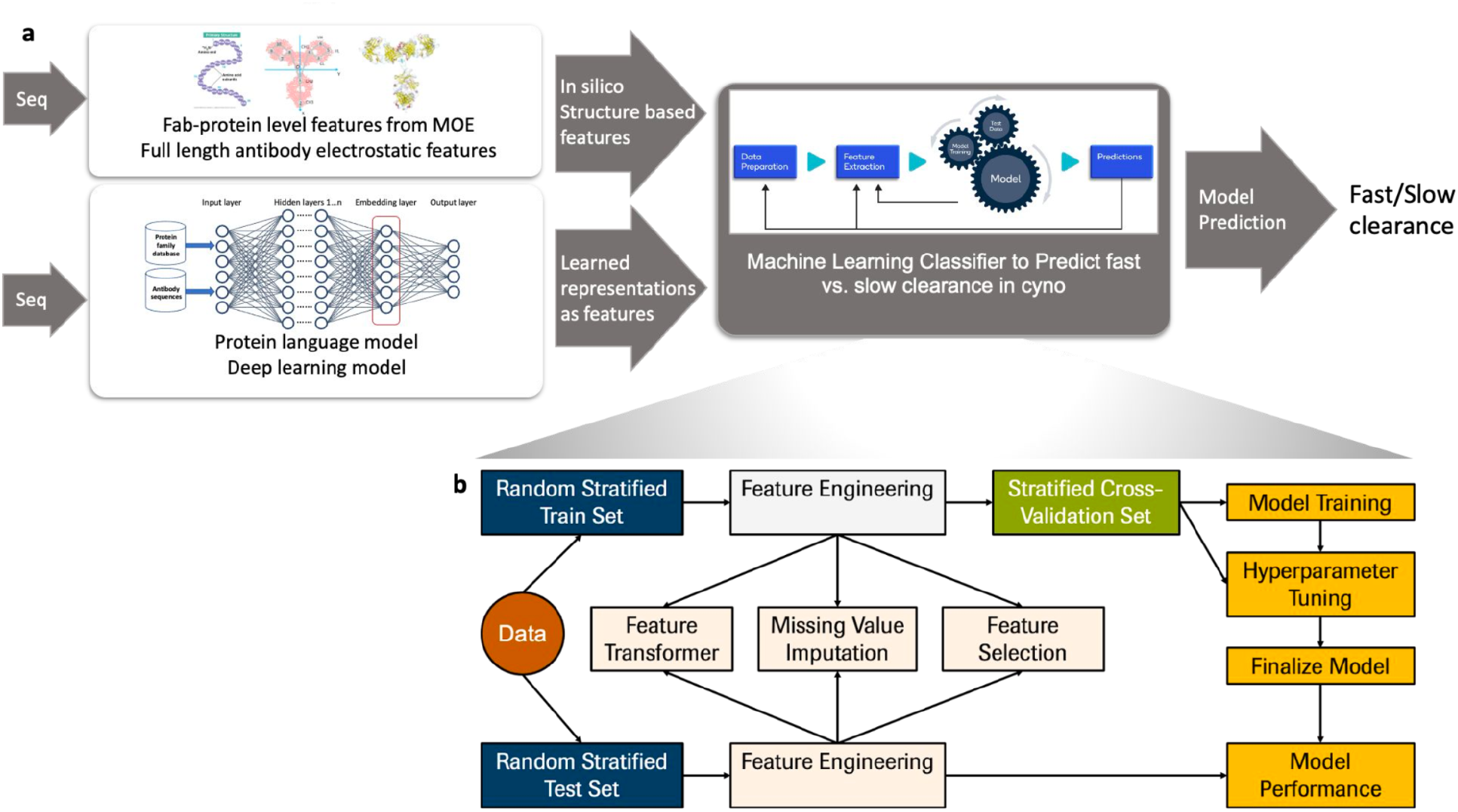
Schematic representation of machine learning model pipeline. (a) overview of how protein language models trained on large datasets can inform machine learning classifiers to predict antibody clearance. (b) Detailed structure of classification pipeline and division of test and training dataset.

**Figure 5.**
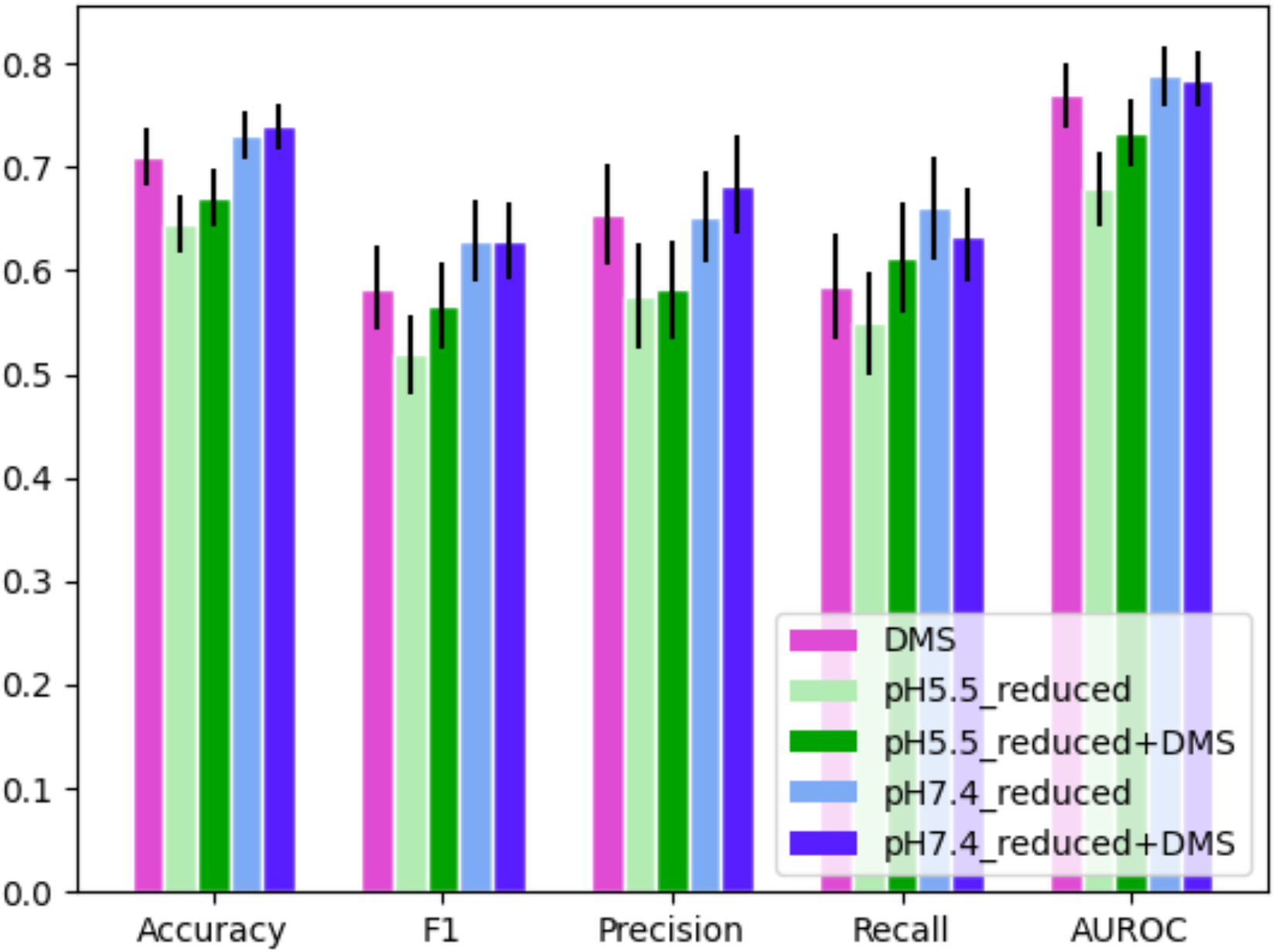
Performance evaluation and comparison of different models. Models trained on the Deep Manifold Sampler (DMS) feature set perform similarly to those trained on the pH 7.4_reduced feature set. Black lines indicate the standard deviation.

Finally, we assessed model performance when both physicochemical features and embeddings were incorporated together. Figure 5 shows the performance comparison of our top classical ML models using only physicochemical features vs models incorporating embeddings in predicting fast versus slow clearance of mAbs in cynos. The pH7.4_reduced+DMS model exhibited the highest numerical performance (accuracy 73.9±1.1%) across all models (Figure 5). There is a slight improvement upon inclusion of the DMS features in precision going from 65.2±2.3% to 68.3±2.4%, representing a reduction in the false discovery rate. The addition of DMS features to the pH5.5_reduced features improved performance from an accuracy of 64.5±1.4% to 67.1±1.4% (Figure 5). Taken together, these results suggest that the embeddings and physicochemical features at pH 7.4 contain primarily overlapping information, enabling comparable performance in classical machine learning models when used individually with little improvement when combined. Both of these feature sets contain more relevant predictive information than the pH 5.5 features, at least for this set of antibodies. For the use case of screening out fast clearing mAbs earlier in the pipeline without eliminating potentially promising normal-clearing mAbs, combining the feature sets to achieve greater precision (i.e. a low false discovery rate) without loss of accuracy is more desirable.

### Feature importance analysis points to the role of electrostatic features in antibody clearance

SHAP analysis of all models was performed to identify the top descriptors influencing the model for prediction of clearance class as shown in Figure 6 and supplementary Figure 5. pH7.4_reduced+DMS shows a combination of physicochemical and embedding features to be important in their decision making process. Features relevant to charge and moment are consistently among the top physicochemical features for these models as well as positive charged patch in the CDR region (Supp. Figure 5). Association of charge with clearance is consistent with what has been previously reported in the literature [3].

**Figure 6.**
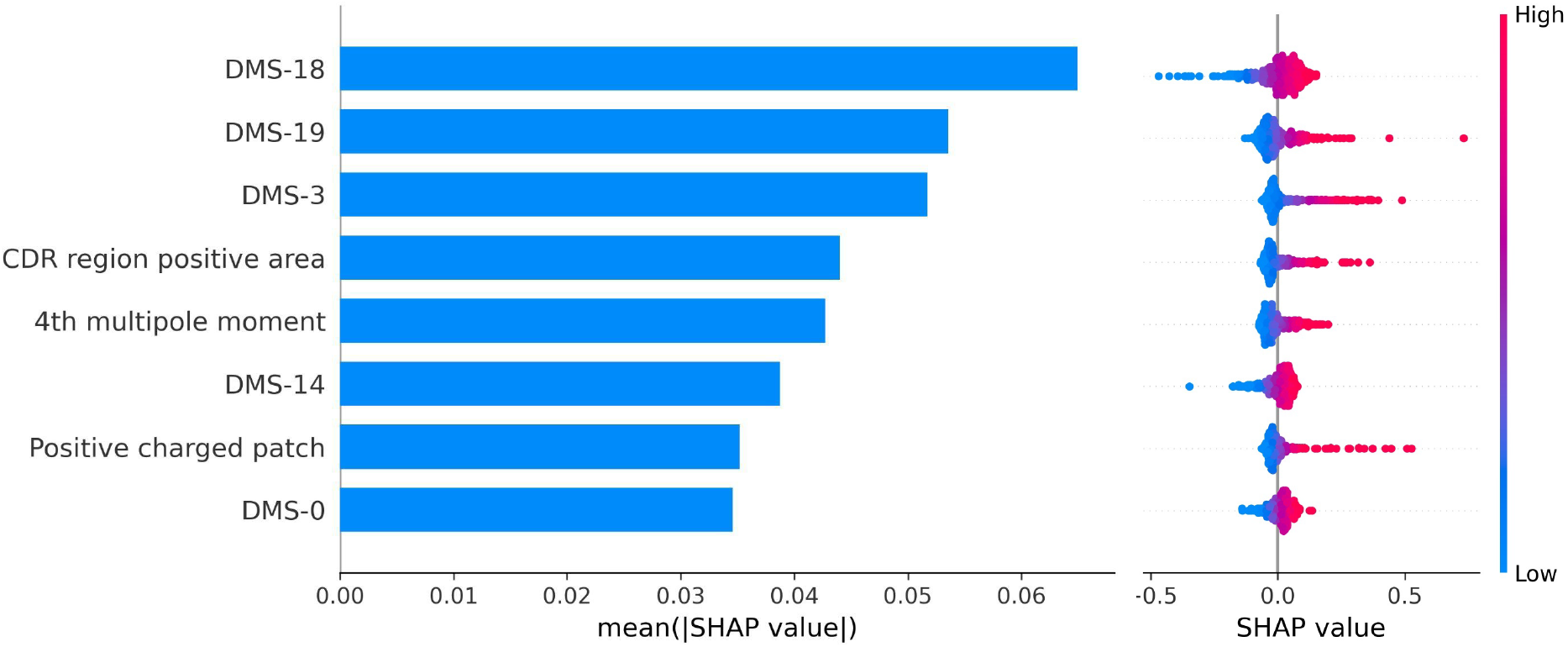
Feature importance for the features of the model trained with pH7.4_reduced+DMS feature sets. Feature importance is reported for the test data. Features of the form “both-n” are from the DMS feature set. Features with charge and moment as well as embeddings are consistently among the most informative features. (Left) Average of absolute impact on model output performance and (Right) dot plot of SHAP value impact on model output performance .

## Discussion

Various technological advancements, including application of artificial intelligence approaches to sequence-based structure and binding prediction, have accelerated the pace and breadth of therapeutic antibody design. As more monoclonal antibody candidates are generated for preclinical and clinical development, the need to consider prioritizing them based in part on feasibility of subsequent development becomes increasingly important [7]. One of the key properties determining the clinical feasibility of mAbs is in vivo clearance. Early detection of mAb candidates with favorable PK and omission of those likely to have fast clearance can reduce time consuming antibody campaigns and subsequent in vitro and animal studies. While assay-based approaches are used for early screens for PK feasibility of mAbs, these still require molecule generation and testing. As more data becomes available for algorithm training, *in silico* approaches such as machine learning predictive approaches become increasingly feasible and promising.

In this work, we developed a machine learning classification model to identify molecules with fast clearance in cynos based only on proposed antibody sequences, using features derived from different *in silico* modeling approaches. While there have been previous reports about sequence based in silico models predicting developability in mice [23], to the best of our knowledge, this is the first time that sequence-based ML and DL models are being used to predict clearance in cynomolgus monkeys. Specifically we have compared the performance of clearance classification based on features from either structure based modeling, protein language models, or a combination of the two. For structural modeling, we considered different subsets of features derived from the MOE environment, with or without feature reduction. For the deep learning approach, we used Deep Manifold Sampler, a protein language algorithm that has been fine-tuned on a dataset of antibodies as opposed to other such models trained on general protein datasets [5,8,10]. Using either modeling approaches for feature generation yielded roughly similar performance (accuracy = 73.1±1.1% vs 71±1.4%, precision = 65.2±2.3% vs 65.5±2.5% respectively). Combining feature reduction and deep-learning based embeddings together yielded accuracy of 73.9±1.1% and precision of 68.3±2.4%, a marginal but highly desirable improvement on precision, lowering false discovery rate. Regardless, the fact that a model trained using only the DMS features achieved similar performance to the physicochemical features is impressive as the DMS features are calculated rapidly and without any tertiary structural information; furthermore, the protein language model based approach does not rely on knowledge-based feature selection or feature reduction. However, an important advantage of the structural modeling based approach is interpretability. We can utilize feature importance metrics like Shapley additive explanations to understand which inputs are driving the models’ predictive capacity. This allows for a ‘sanity check’ to see if the important features seem physically relevant, provides input into molecular characteristics and processes influencing clearance, and informs the inclusion of new types of features in the future. The structural models are not designed for PK prediction, although manual feature engineering leveraging hypotheses of structural property combinations that influence PK could further improve performance. Conversely, since the current DMS embeddings are fine-tuned on unlabeled data in a self-supervised method, additional improvements might be achieved by fine tuning the deep learning model on labeled dataset of fast vs slow clearance, as well as with other developability criteria. As datasets of antibodies grow, deep learning models trained specifically on antibodies can be used to predict PK properties like clearance. Finally, additional training data will be key to improving the model, as the relatively small dataset limits the performance achievable.

To our knowledge this is the largest dataset compiled of monoclonal antibodies with measured clearance in cyno; however, data availability is still the key limiting factor of our analysis. Machine learning and specifically deep learning models require large datasets to achieve confidence in their predictions and avoid overfitting to limited sets of observations. Therefore, a clear direction for improvement in the models would be to increase the dataset size. We also note a small bias (60.2% slow vs 39.8% fast) in our dataset. Molecules that progress to testing in cynos have typically already passed early in vitro screening assays and have undergone PK testing in rodent species where molecules showing signs of fast clearance are often eliminated from further consideration. Reiterating that early adoption of sequence based approaches can reduce the number of animal experiments.

We incorporated several approaches to compensate for the limited dataset size. First, we developed classical machine learning methods which are not as data hungry as deep learning approaches. By using feature reduction techniques we were able to improve accuracy from 67.7±1.2% to 73.1±1.1% with a precision of 65.2±2.3% (model pH7.4_reduced Fig 3), while maintaining interpretability. In particular, hydrophobicity and charge were highlighted by importance analysis as influential features, consistent with previously published results [3](Supp. Fig 5). Physicochemical features calculated for pH 5.5 yielded poorly performing models and when combined with the other feature sets, did not improve performance relative to models based on pH 7.4 or DMS embeddings alone. These results suggest that, at least for this set of conventional (non-Fc engineered) mAbs, not only physicochemical features at pH 5.5 are not good predictors of non-specific clearance, they do not provide any complementary information to the features at neutral pH either. In fact, inclusion of pH5.5 features (model pH5.5_reduced+pH7.4_reduced; model pH5.5_reduced+DMS) performed slightly worse than models based on pH7.4 or DMS embeddings alone. Notably, however, we did not use expert-knowledge informed feature reduction or combination approaches. While these might theoretically improve performance, we sought objective, automated approaches to modeling for generalizability and efficiency.

Second, we found that we needed a nested cross-validation approach in order to properly estimate model performance. Generally, a test set is held back from the training process to use for a final estimate of a model’s performance. This out-of-sample approach provides the best estimate of how the model will perform on the truly new data it will encounter after deployment. Unfortunately, owing to the small dataset size, our test sets were on the order of 20 molecules and thus were highly variable in their composition - some molecules are simply more predictable than others. This meant that the test set performance was determined more by the constituents of the test set than the quality of the model. We mitigated this issue by repeating the model training and testing procedure using Monte Carlo cross validation and averaging the results.

In this work, we focused on the classification task of predicting fast versus slow clearance in cyno based on a cut-off of 8 mL/kg/day. While other cut-offs like 10 mL/kg/day have been previously used in the literature [3], a more recent analysis of scaling cyno to human clearance supports 8 mL/kg/day as a useful threshold to identify mAbs with greatest likelihood of meeting common dosing regimens [19]. A possible improvement on this current work would be to predict the actual clearance value which is developing a regression as opposed to a classification model. However, development of a regression model with acceptable performance would require a much larger dataset with a more uniform distribution of clearance values. Our current dataset is not only small in size, but has outliers with clearance values as large as >100 mL/kg/day which makes the regression task increasingly difficult.

One of the key benefits of our in-silico proposed method is its generalizability. Cyno clearance is one of the best predictors of human clearance of mAbs [11] and gathering sufficient data on human molecules is much more challenging. As more data on other kinds of molecules - such as engineered antibodies that include FcRn modifications - become available, the methods presented here can be applied to these datasets. Given a sufficiently large dataset with other measured PK characteristics such as volumes of distribution, half-lives and immunogenicity, ML/DL methods can be developed to predict these characteristics using similar approaches discussed in this work. Moreover, multi-task learning approaches can be leveraged that allows for simultaneous prediction of these characteristics to inform the broader developability of molecules very early on in the pipeline of drug development. Finally, for later stages of research and development, data from screening assays and early preclinical studies can be incorporated in these models or in the screening process to further improve lead selection and PK translation.

Ultimately, we believe this work lays the groundwork and provides proof of concept for a significant advance in antibody PK prediction that leverages the recent revolution in sequence-based deep learning for proteins. Our results further highlight the relevance of sequence-based predictions for assessment of therapeutic developability criteria. Future training of these algorithms on PK, immunogenicity, and other developability criteria could revolutionize discovery and development of therapeutic antibodies and proteins in general. Finally, these results point to the broader promise of transfer learning applications of protein language models of structure and binding to address critical questions in protein therapeutic development.

## Acknowledgements

We would like to thank Hao Cai, Douglas Leipold, Radhika Kenkre for their help in dataset curation. We would also like to thank Dan Berenberg, Stephen Ra and Richard Bonneau for providing access to the deep manifold sampler embedding pipeline. The authors thank Greg Ferl, Hao Cai and Isidro Hotzel as well for their insightful comments on the manuscript.

## Materials and Methods

### Homology modeling and physicochemical feature calculations

The software Molecular Operating Environment (MOE) [11] from Chemical Computing Group was used to build the homology models of fragment antigen-binding (Fab) domains. The force field was set to the default Amber10 EHT using an internal and external dielectric values of 4 and 80, the non-bonded cutoff distances of 10 and 12 Å, and the Born solvation model. The prepared structures were energy minimized to the root mean square gradient (RMSG) below 0.00001 kcal/mol/Å2.

Protein properties for each Fab region were calculated using the Ensemble pH sampling module available in MOE 2019 [11], spanning two pH ranges, 4.5 to 6.5 (targeting pH=5.5) and 6.4 to 8.4 (targeting pH=7.4). The number of conformations per unit of pH range was set to 50. The protein properties were calculated as Boltzmann weighted averages of the individual values from an ensemble of 100 total conformations.

In addition to protein property features computed in MOE, we also calculated the multipole moments and the coarse-grained charges of the full-length antibody structures using the following protocol. The full-length antibodies were produced by aligning (alpha-carbon root-mean square deviation minimization) the Fab models to either the left or right arm of a generic full-length IgG1 molecule. The mab IgG1 framework was based on a full structure of the IgG1 antibody generated by Brandt et al. [17] and equilibrated to a relaxed conformation. The C-terminal lysines were cleaved off from the IgG1 homology models. The constant domain (Fc) and Fab arms in this mab IgG1 framework were oriented so that the Fc is aligned with the x axis and the Fab arms are extended toward the y axis (as shown in Figure 2). The full-length homology models were energy minimized using SANDER, available in AMBER 2015 (32). PDB2PQR (33) (v. 1.8) was used with the AMBER force field (34) to assign partial atomic charges whereby ionization states were determined using PROPKA (35,36) (pH 5.5 and 7.4).

The partial atomic charges on the full-length antibody structure were used to calculate the low-order molecular multipole moments (m1-m6) and the coarse-grained charges (c1-c6) in the expansion of the full-length antibody charge distributions using an in-house script. The mathematical equations used to calculate the multipole moments and the coarse grained charges are provided in Ref. [13].

### Machine learning models

We optimized model architecture and hyperparameters to find the algorithm that performed best for our particular problem using Tree-Based Pipeline Optimization Tool (TPOT), an AutoML approach. For each set of features, 50,000 model pipelines were generated by the TPOT for Python [20]. Each pipeline randomly sampled from a set of classifier algorithms, selectors, preprocessing steps, and associated hyperparameters. A table with the available components and hyperparameters can be found in Supplementary Table 2. Pipelines were evaluated by Monte Carlo bootstrapping. In this procedure, the dataset was repeatedly split into training, validation, and test sets (in an 80-10-10 ratio). The pipeline was fit on the training set, and evaluated on the validation and test sets. The performance metrics were averaged over 100 random repeated splits. The pipeline which performed best on average over the validation sets was selected. Its out-of-sample performance was estimated from the average test set performance, which is the value reported in this paper.

The feature set consisted of a number of highly correlated features (Supp. Figure 3). This forced pipelines to sift through redundant and non-informative features hindering efficient training. Although the generated pipelines could include a feature selection layer like recursive feature elimination, this was not a requirement and was computationally expensive to run. We selected a subset of features to use (termed “reduced”) by examining which features were routinely calculated as important by permutation feature importance. For this calculation, we selected the top 20 performing models for a particular dataset. These models were repeatedly trained and applied to random training splits with permutation feature importance being calculated on the corresponding test splits. These feature importances were averaged over all models and the top 15 features were selected for the reduced set. In the case of pH7.4_reduced, only 13 features had non-zero importance and so that feature set consists of only those features.

To reduce the hyperparameter search space, we narrowed our pool of classifier algorithms down to decision tree, gradient boost, and Gaussian naive Bayes. This set was determined through preliminary experimentation in which it was found that these algorithms were typically more effective than the alternatives.

We generated sets of sequence-based features from a pre-trained protein language model. The model investigated, Deep Manifold Sampler, was first developed by Gligorijevic et al. [8] and further fine tuned on antibodies by Berenberg et al. [10] . This model was trained on the diverse proteins of the Pfam dataset and further fine tuned on native human antibodies from OAS. This model yielded a 256 feature embedding for each antibody. The heavy and light chains of the antibodies were embedded together. The whole-protein embedding was produced by averaging the individual amino acid embeddings. These features contained some redundancy for our dataset and so principal components analysis (PCA) on the entire dataset was used for feature reduction. An arbitrary cutoff of the first 20 components from PCA were selected. These features accounted for 98% of the variance observed in the DMS embeddings. These 20 features were concatenated to the pH5.5_reduced and pH7.4_reduced feature sets to create hybrid feature sets for model training. The similarity between certain DMS features and the pH 7.4 physicochemical features is shown in Supplementary Figure 4.

