## Supplementary figures and tables for "Leveraging protein language and structural models for early prediction of antibodies with fast clearance"

Supplementary Figure 1
Box plot association of variable domain and clearance

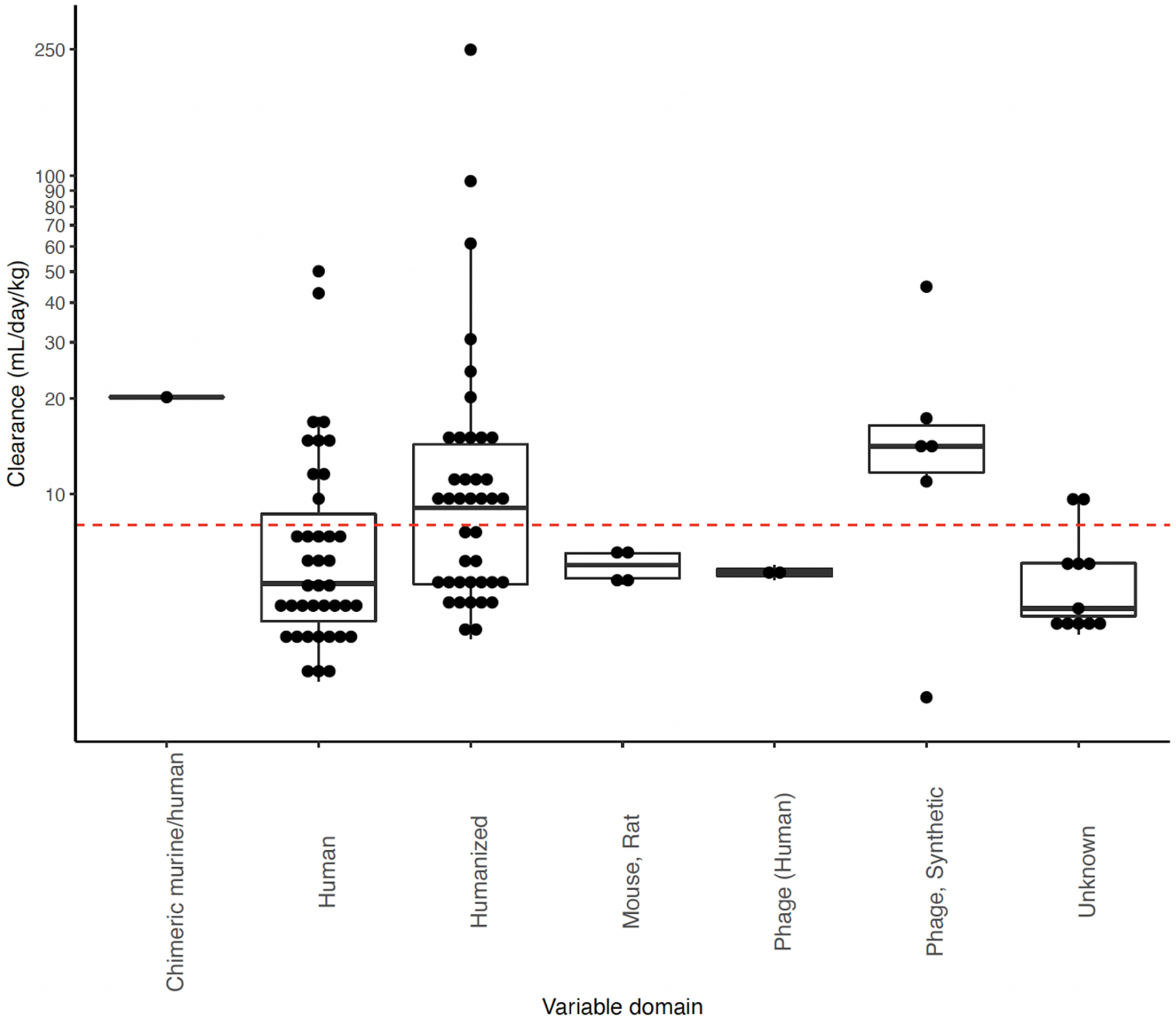


Supplementary Figure 2. Wilcoxon test assessment shows only 4 out of 33 physicochemical features are significantly different between slow and fast molecules. Significant is assigned based on p-value < 0.05. Significant associations are colored yellow.

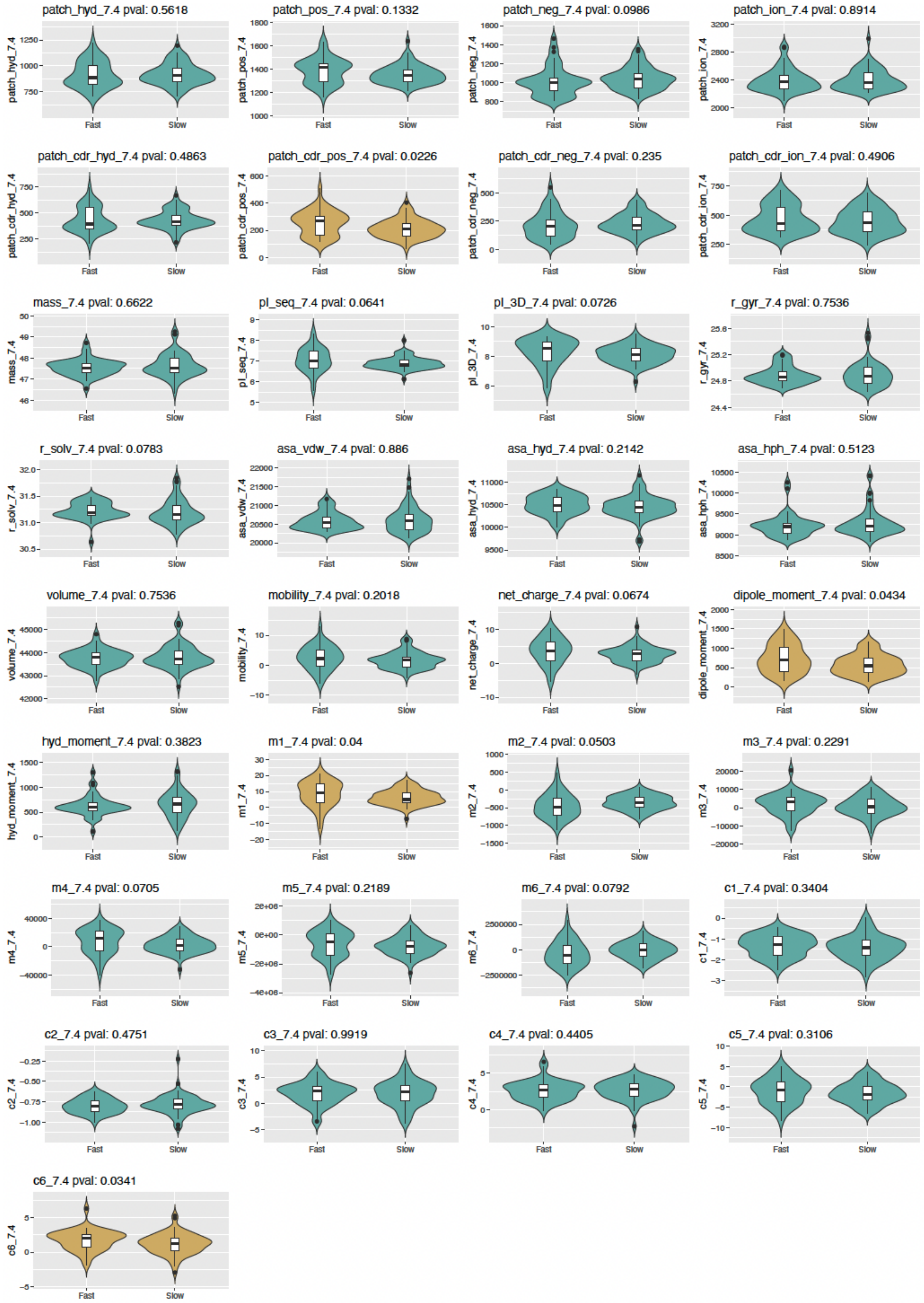

patch_hyd_7.4 pval: 0.5618
patch_pos_7.4 pval: 0.1332
patch_neg_7.4 pval: 0.0986
patch_ion_7.4 pval: 0.8914
patch_cdr_hyd_7.4 pval: 0.4863
patch_cdr_pos_7.4 pval: 0.0226
patch_cdr_neg_7.4 pval: 0.235
patch_cdr_ion_7.4 pval: 0.4906
mass_7.4 pval: 0.6622
pI_seq_7.4 pval: 0.0641
pI_3D_7.4 pval: 0.0726
r_gyr_7.4 pval: 0.7536
r_solv_7.4 pval: 0.0783
asa_vdw_7.4 pval: 0.886
asa_hyd_7.4 pval: 0.2142
asa_hph_7.4 pval: 0.5123
volume_7.4 pval: 0.7536
mobility_7.4 pval: 0.2018
net_charge_7.4 pval: 0.0674
dipole_moment_7.4 pval: 0.0434
hyd_moment_7.4 pval: 0.3823
m1_7.4 pval: 0.04
m2_7.4 pval: 0.0503
m3_7.4 pval: 0.2291
m4_7.4 pval: 0.0705
m5_7.4 pval: 0.2189
m6_7.4 pval: 0.0792
c1_7.4 pval: 0.3404
c2_7.4 pval: 0.4751
c3_7.4 pval: 0.9919
c4_7.4 pval: 0.4405
c5_7.4 pval: 0.3106
c6_7.4 pval: 0.0341
Fast
Slow

Supplementary Figure 3. Pearson correlation coefficient for physicochemical features at pH 7.4.

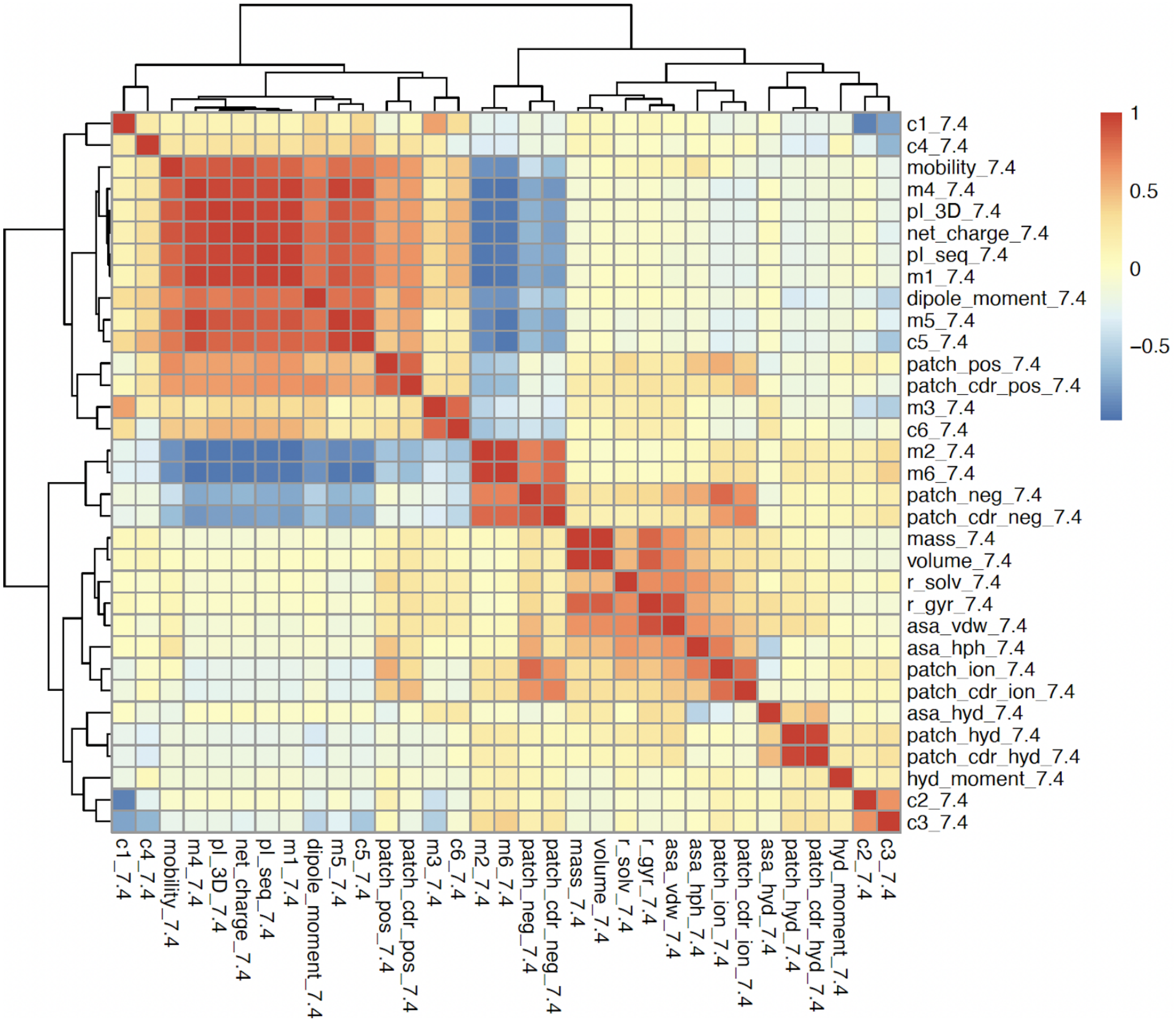


Supplementary Figure 4. Pearson correlation coefficients for pH 7.4 and DMS features.

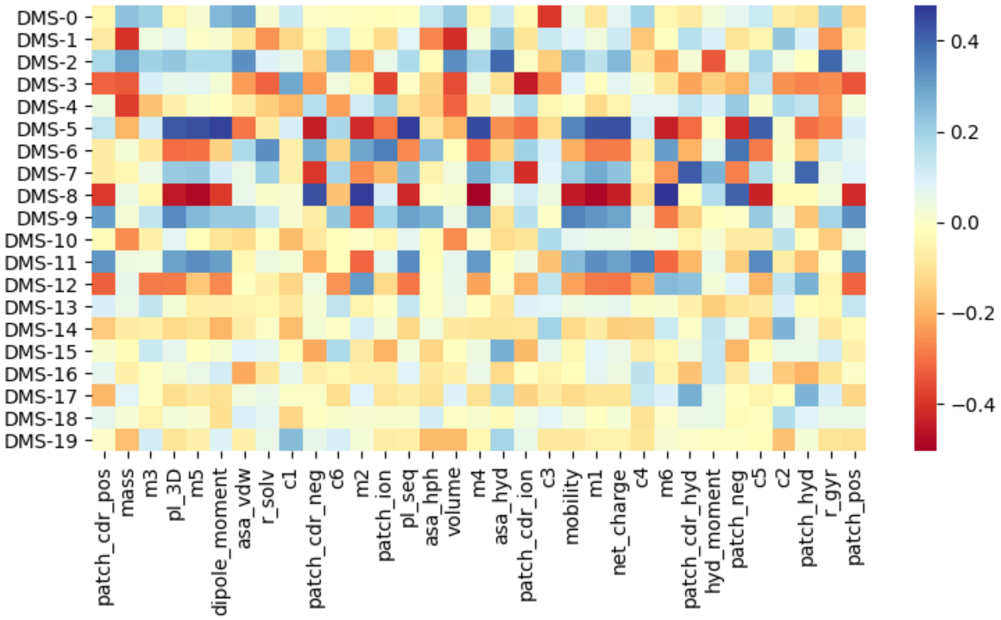


Supplementary Figure 5. Feature importance for the features of the model trained with various feature sets. Feature importance is reported for the test data. Features of the form “DMS-n” are from the DMS feature set. Structural features are calculated at pH 7.4 unless otherwise noted. Features with charge and moment as well as embeddings are consistently among the most informative features. Dot plots of SHAP values are shown on the left and average absolute impact on the right for (a) pH5.5, (b) pH5.5_reduced, (c) pH5.5_reduced+DMS, (d) pH7.4, (e) pH7.4_reduced, (f) pH7.4_reduced+DMS, (g) pH5.5_reduced+pH7.4_reduced, and (h) DMS.

(a)

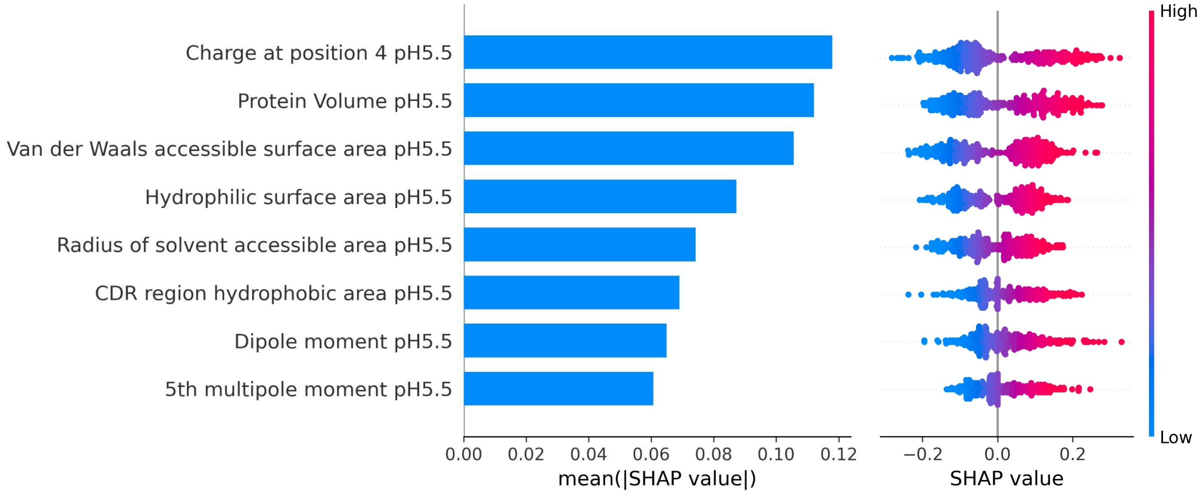


(b)

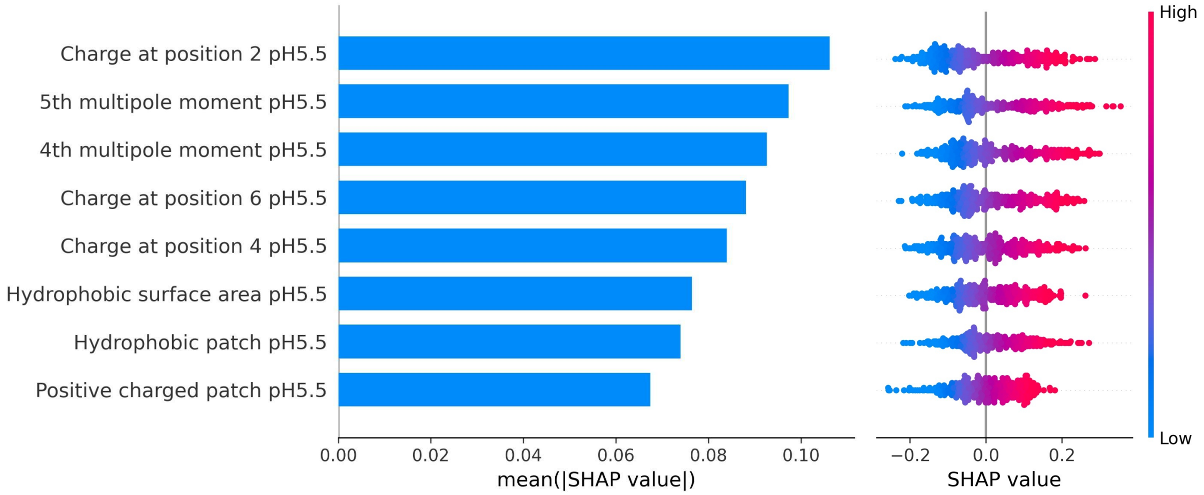

Charge at position 2 pH5.5
5th multipole moment pH5.5
4th multipole moment pH5.5
Charge at position 6 pH5.5
Charge at position 4 pH5.5
Hydrophobic surface area pH5.5
Hydrophobic patch pH5.5
Positive charged patch pH5.5
0.00 0.02 0.04 0.06 0.08 0.10
mean(|SHAP value|)
−0.2 0.0 0.2
SHAP value
High
Low


(c)

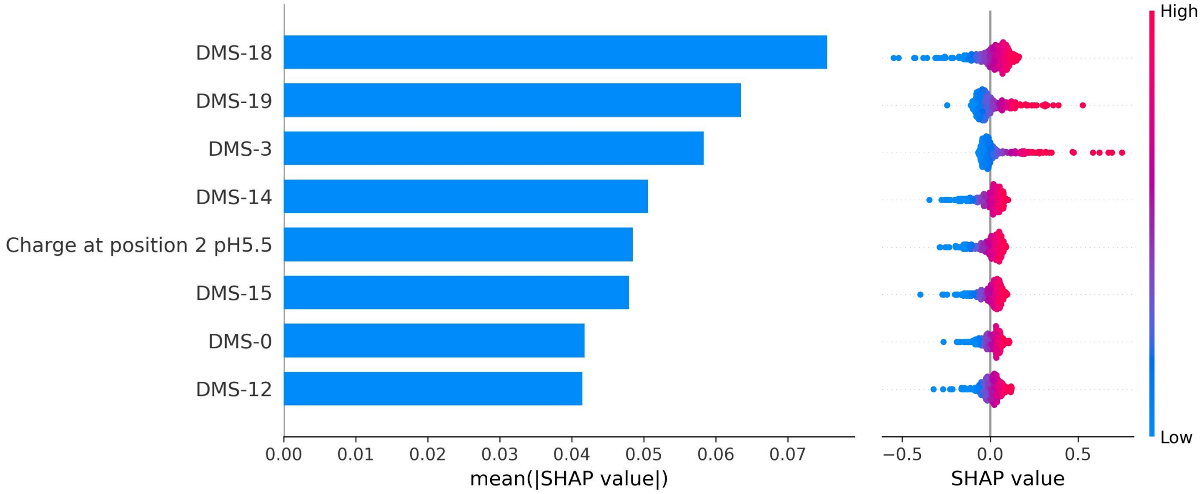

DMS-18
DMS-19
DMS-3
DMS-14
Charge at position 2 pH5.5
DMS-15
DMS-0
DMS-12
0.00 0.01 0.02 0.03 0.04 0.05 0.06 0.07
mean(|SHAP value|)
−0.5 0.0 0.5
SHAP value
High
Low


(d)

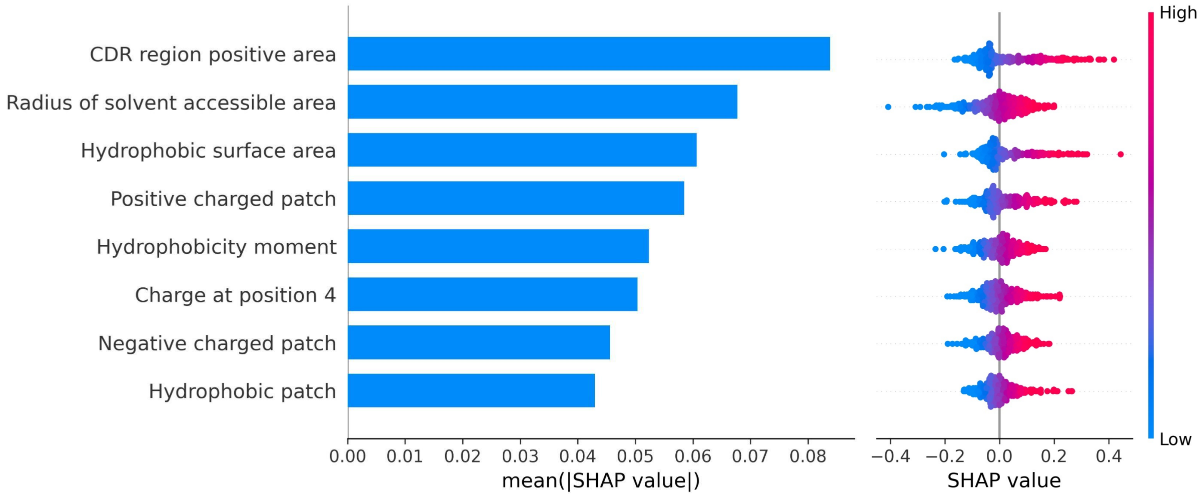

CDR region positive area
Radius of solvent accessible area
Hydrophobic surface area
Positive charged patch
Hydrophobicity moment
Charge at position 4
Negative charged patch
Hydrophobic patch
0.00 0.01 0.02 0.03 0.04 0.05 0.06 0.07 0.08
mean(|SHAP value|)
−0.4 −0.2 0.0 0.2 0.4
SHAP value
High
Low


(e)

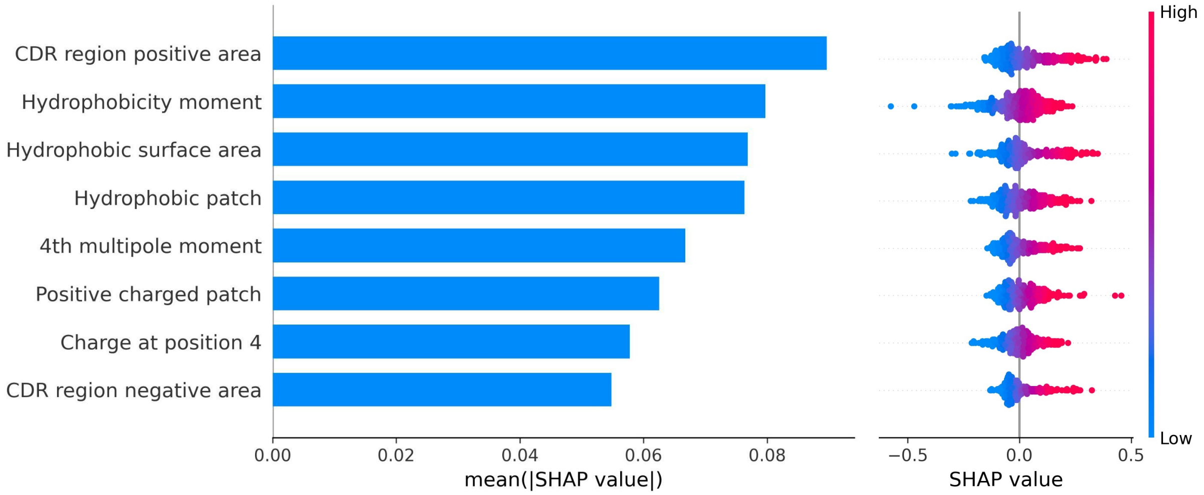
CDR region positive area
Hydrophobicity moment
Hydrophobic surface area
Hydrophobic patch
4th multipole moment
Positive charged patch
Charge at position 4
CDR region negative area
0.00
0.02
0.04
0.06
0.08
mean(|SHAP value|)
−0.5
0.0
0.5
SHAP value
High
Low

(f)

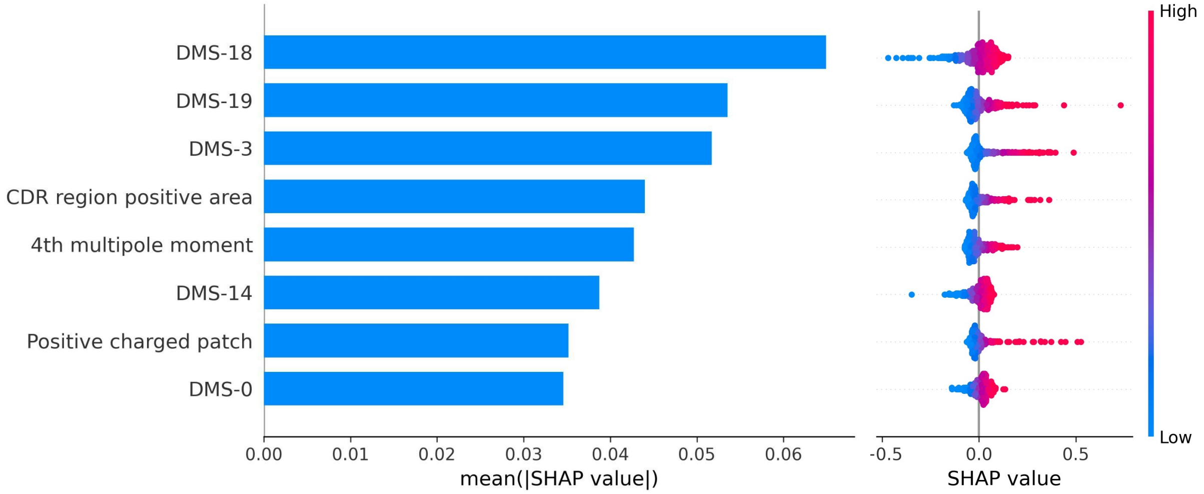
DMS-18
DMS-19
DMS-3
CDR region positive area
4th multipole moment
DMS-14
Positive charged patch
DMS-0
0.00
0.01
0.02
0.03
0.04
0.05
0.06
mean(|SHAP value|)
-0.5
0.0
0.5
SHAP value
High
Low

(g)

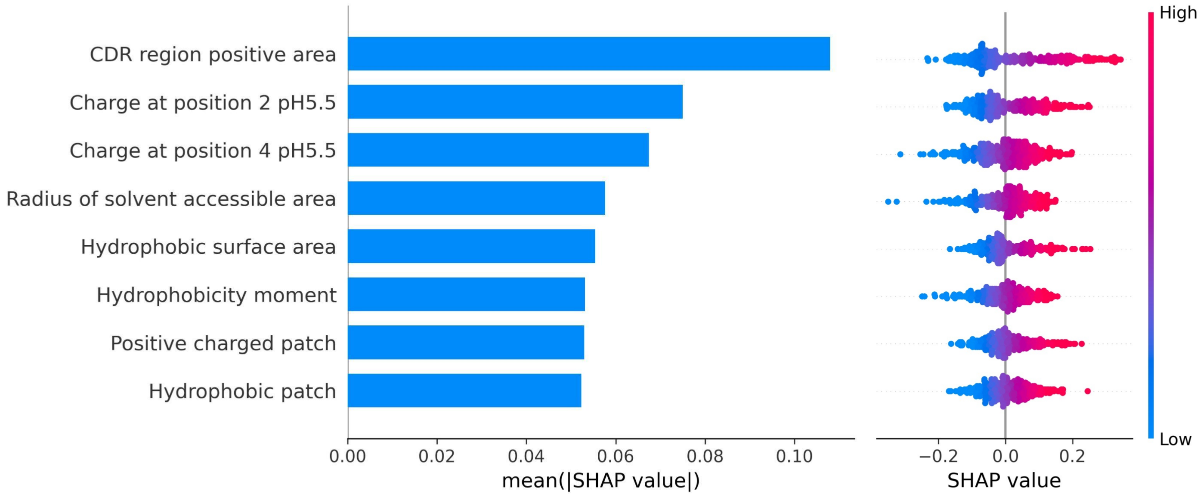
CDR region positive area
Charge at position 2 pH5.5
Charge at position 4 pH5.5
Radius of solvent accessible area
Hydrophobic surface area
Hydrophobicity moment
Positive charged patch
Hydrophobic patch
0.00
0.02
0.04
0.06
0.08
0.10
mean(|SHAP value|)
−0.2
0.0
0.2
SHAP value
High
Low

(h)

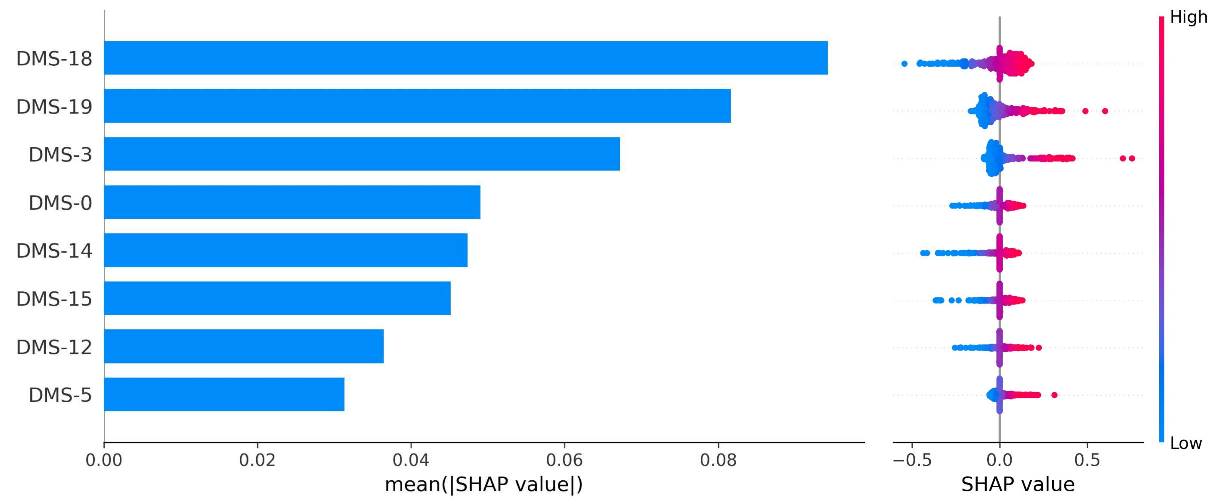


Supplementary Table 1. List of physicochemical features included in pH 5.5 and pH 7.4 sets

| | Feature name | Description |
|---|---|---|
| 1 | patch_hyd | Hydrophobic patch |
| 2 | patch_pos | Positive charged patch |
| 3 | patch_neg | Negative charged patch |
| 4 | patch_ion | Ionized charged patch |
| 5 | patch_cdr_hyd | Hydrophobic patch in CDR region |
| 6 | patch_cdr_pos | Positive charged patch in CDR region |
| 7 | patch_cdr_neg | Negative charged patch in CDR region |
| 8 | patch_cdr_ion | Ionized charged patch in CDR region |
| 9 | mass | Molecular weight |
| 10 | pI_seq | pI based on AA sequence |
| 11 | pI_3D | pI based on 3D structure |
| 12 | r_gyr | Radius of gyration |
| 13 | r_solv | Radius of solvent accessible area |
| 14 | asa_vdw | Accessible surface area, van der Waals |
| 15 | asa_hyd | Accessible surface area, hydrophobic |
| 16 | asa_hph | Accessible surface area, hydrophilic |
| 17 | volume | Protein volume |
| 18 | mobility | Protein Mobility |

| | | |
|---|---|---|
| 19 | net_charge | Net charge based on the sequence |
| 20 | dipole_moment | Protein dipole moment by MOE |
| 21 | hyd_moment | Hydrophobicity moment |
| 22 | m1 | Monopole moment (net charge) |
| 23 | m2 | Dipole expansion moments |
| 24 | m3 | Advanced expansion moments |
| 25 | m4 | Advanced expansion moments |
| 26 | m5 | Advanced expansion moments |
| 27 | m6 | Advanced expansion moments |
| 28 | c1 | Charge at position 1 |
| 29 | c2 | Charge at position 2 |
| 30 | c3 | Charge at position 3 |
| 31 | c4 | Charge at position 4 |
| 32 | c5 | Charge at position 5 |
| 33 | c6 | Charge at position 6 |

Supplementary Table 2. List of pipeline components and the range of hyperparameter values used in optimization. The components are standard pieces from the Scikit-learn Python library. Classifiers are colored green, preprocessors are colored blue, and selectors are colored red.

| **Component** | **Hyperparameter** | **Values** |
|---|---|---|
| ExtraTreesClassifier | criterion | [‘gini’, ‘entropy’] |
| | n_estimators | [10, 15, 20, 25, 50, 100] |
| | min_samples_split | range(2, 21) |
| | min_samples_leaf | range(1, 21) |
| | bootstrap | [True, False] |
| | max_features | numpy.arange(0.05, 1.01, 0.05) |
| GradientBoostingClassifier | n_estimators | [10, 15, 20, 25, 50, 100] |
| | learning_rate | [1e-3, 1e-2, 1e-1, 0.5, 1.] |
| | max_depth | range(1, 11) |

| | | |
|---|---|---|
| | min_samples_split | range(2, 21) |
| | min_samples_leaf | range(1, 21) |
| | subsample | numpy.arange(0,05, 1.01, 0.05) |
| | max_features | numpy.arange(0.05, 1.01, 0.05) |
| GaussianNB | - | - |
| Binarizer | threshold | numpy.arange(0.0, 1.01, 0.05) |
| FastICA | tol | numpy.arange(0.0, 1.01, 0.05) |
| FeatureAgglomeration | linkage | [‘ward’, ‘complete’, ‘average’] |
| | affinity | [‘euclidean’, ‘l1’, ‘l2’, ‘manhattan’, ‘cosine’] |
| MaxAbsScaler | - | - |
| MinMaxScaler | - | - |
| Normalizer | norm | [‘l1’, ‘l2’, ‘max’] |
| Nystroem | kernel | [‘rbf’, ‘cosine’, ‘chi2’, ‘laplacian’, ‘polynomial’, ‘poly’, ‘linear’, ‘additive_chi2’, ‘sigmoid’] |
| | gamma | numpy.arange(0.0, 1.01, 0.05) |
| | n_components | range(1, 11) |
| PCA | svd_solver | [‘randomized’] |
| | iterated_power | range(1, 11) |
| RBFSampler | gamma | numpy.arange(0.0, 1.01, 0.05) |
| RobustScaler | - | - |
| StandardScaler | - | - |
| OneHotEncoder | min_frequency | [0.05, 0.1, 0.15, 0.2, 0.25] |
| | sparse | [False] |

| | | |
|---|---|---|
| SelectFwe | alpha | numpy.arange(0, 0.05, 0.001) |
| | score_func | {‘sklearn.feature_selection.f_classif’: None} |
| SelectPercentile | percentile | range(1, 100) |
| | score_func | {‘sklearn.feature_selection.f_classif’: None} |
| VarianceThreshold | threshold | [0.0001, 0.0005, 0.001, 0.005, 0.01, 0.05, 0.1, 0.2] |
| RFE | step | numpy.arange(0.05, 1.01, 0.05) |
| | estimator | 'sklearn.ensemble.ExtraTreesClassifier': {'n_estimators': [100], 'criterion': ['gini', 'entropy'], 'max_features': np.arange(0.05, 1.01, 0.05) |
| SelectFromModel | threshold | numpy.arange(0, 1.01, 0.05) |
| | estimator | 'sklearn.ensemble.ExtraTreesClassifier': {'n_estimators': [100], 'criterion': ['gini', 'entropy'], 'max_features': np.arange(0.05, 1.01, 0.05) |
